# Oscillatory dynamics underlying emotion-cognition integration: differential role of theta and alpha oscillations

**DOI:** 10.1101/2022.03.21.485174

**Authors:** Zhihao Wang, Katharina S. Goerlich, Mai Chen, Pengfei Xu, Yuejia Luo, André Aleman

## Abstract

A fundamental aspect of human mental life is the seamless ability for integration of emotion and cognition. Despite progress regarding the spatial architecture of Emotion-Cognition Integration (ECI), the time course of ECI processes remains unclear. To examine the temporal organization of brain oscillations underpinning ECI, we simultaneously manipulated emotional valence of stimuli and cognitive task demand while recording electrophysiological responses of 61 participants. They were asked to complete tasks with low (body-part judgement) and high (laterality judgement) cognitive demand while viewing other people photographs that varied on dimensions of laterality (left or right), body-part (hand or foot), and emotional valence (pain or no pain). We found increased reaction times and error rates in pain versus no pain during laterality judgement relative to body-part judgement, suggesting reciprocal inhibition between emotion and cognition. EEG results showed that 1) emotion processing (valence) occurred first in the theta band from 144 to 372 ms; 2) cognitive processing (laterality) took place in the theta band from 332 to 608 ms; 3) emotional and cognitive processes were integrated in the alpha band from 268 ms and lasted to 800 ms. These findings reveal oscillatory dynamics of the processing and integration of emotion and cognition, providing further insights into the underlying neurophysiology. This may ultimately contribute to our understanding of ECI processing in psychopathology.

## Introduction

A fundamental aspect of human mental life is the seamless ability for integration of emotion and cognition. The historical view of “functional specialization” holds that different brain areas are specialized for emotion and cognition. Inspired by this viewpoint, many studies examined either the influence of cognition on the “emotional brain” (Ochsner and Gross, 2005) or the impact of emotion on the “cognitive brain” (Dolan, 2002; Pessoa, 2009). However, the concept of functional localization alone is not sufficient to explain the highly complex and dynamic nature of human behavior (Pessoa, 2008). Indeed, in recent years research emphasizes functional integration of brain networks in order to process emotional and cognitive information (Okon-Singer *et al.*, 2015). Emotion-cognition integration (ECI) occurring at both the behavioral and neural levels are of great importance for adaptation to one’s environment and thus to successful everyday functioning (Pessoa, 2008). Specifically, emotion and cognition not only strongly interact in the brain but are dynamically integrated to jointly contribute to behavior.

Gu et al., (2013) investigated the interaction between cognitive and affective processes using an empathy-for-pain paradigm that simultaneously manipulated cognitive task demand and affective valence of visual stimuli. Using functional Magnetic Resonance Imaging (fMRI), they identified an interaction between cognitive demand and affective stimulus valence in the anterior insular cortex as well as in the primary somatosensory cortex and regions of the frontoparietal network. This may be indicative of a role as interface for ECI in these regions. However, the temporal resolution of MRI is low, and thus the temporal dynamics underpinning ECI remain elusive. Understanding how ECI processes unfold over time is of crucial importance for affective and cognitive neuroscience, given that our brain works dynamically across multiple timescales (Olivers, 2007; Hutchison *et al.*, 2013; King and Dehaene, 2014).

Electroencephalogram (EEG) techniques have excellent temporal resolution in the range of milliseconds and are thus well-suited to shed light on the time course of ECI (Cohen, 2014). Posterior alpha oscillations have been associated with inhibition processes (Klimesch *et al.*, 2007; Jensen and Mazaheri, 2010; Klimesch, 2012), with increased alpha reflecting inhibition and decreased alpha the release from inhibition (Klimesch, 2012; Van Diepen *et al.*, 2019). Given the inhibitory nature of the relationship between emotion and cognition, alpha power can thus be used to examine neurophysiological processes underlying ECI. In addition, frontal-central theta oscillations are of importance for both emotional and cognitive processing (Cohen *et al.*, 2007; Knyazev *et al.*, 2009; Nigbur *et al.*, 2011; Cavanagh and Frank, 2014; Puma *et al.*, 2018). Specifically, decreased theta power has been shown in negative as compared to positive valence processing (Kamarajan *et al.*, 2008), whereas stronger theta power has been observed for high relative to low cognitive demand (Cavanagh and Frank, 2014). In particular, theta power may be a compelling candidate mechanism for such integrative operation given the important role of theta power in cognitive regulation of negative emotion (Ertl *et al.*, 2013).

Here, we tested oscillatory dynamics underlying ECI while recording EEG signals during an empathy-for-pain task adapted from Gu and colleagues, in which we simultaneously manipulated cognitive task demand and affective valence of the stimuli (Gu *et al.*, 2013). First, we replicated ECI at the behavioral level (i.e., reaction times and error rates). Second, a data-driven method was used to examine the temporal organization of different information processing stages. Based on the dual-system model proposing a fast emotional system and a slow cognitive system during human decision making (Kahneman, 2003; Kuo *et al.*, 2009), we predicted that emotion processing, as indexed by theta power, would occur first. Importantly, we predicted an interaction effect between emotion and cognition in theta and alpha oscillations. We also took into account individual differences in propensity for emotion processing and regulation, as this may be a factor of interest. More specifically, this regards alexithymia, which refers to a reduced ability to identify, describe and regulate one’s feelings (Luminet *et al.*, 2018). Given cognitive and emotional deficits in alexithymia (for a review, see Luminet *et al.*, 2021), we explored associations between alexithymia levels and ECI ability.

## Methods and Materials

### Participants

Sixty-four right-handed healthy adults participated in the experiment. Based on the Chinese version of the 20-item Toronto Alexithymia Scale (TAS-20; Bagby et al., 1994) and in light of the international cutoff to access alexithymia with the TAS-20 (Taylor *et al.*, 1988), 32 participants with TAS-20 scores higher or equal to 61 (13.81% of the pool of 695 students; high alexithymia group; HA) and 32 participants with TAS-20 scores lower or equal to 51 (53.24% of the pool; low alexithymia group; LA) participated in the experiment (for demographic information, see Table S1). This sampling procedure ensured sufficient variability in TAS-20 scores. The current sample size was similar to previous studies examining emotion processing in alexithymia, which showed strong effects (e.g., Roedema and Simons, 1999; Heinzel *et al.*, 2010). Data of three participants were excluded from analyses because of excessive EEG artifacts. The final sample consisted of 61 participants (32 HAs; 32 females; age: 17 – 28 years, mean ± SD: 20.49 ± 2.23 years). All participants had normal or corrected-to-normal vision and reported no psychiatric illness in present or past. This study was approved by the Ethics Committee of Shenzhen University and informed written consent was obtained from all participants.

### Self-report questionnaires

In addition to the TAS-20, participants also completed the Chinese version of the Bermond-Vorst Alexithymia Questionnaire (BVAQ; Wang et al., 2021), the Beck Anxiety Inventory (BAI; Beck et al., 1988), the Beck Depression Inventory (BDI; Beck, 1967) and the Autism Spectrum Quotient (ASQ; Baron-Cohen et al., 2001) given that anxiety, depression, and autism have been demonstrated to co-occur with alexithymia (Hendryx *et al.*, 1991; Bird and Cook, 2013; Li *et al.*, 2015).

### Picture Stimuli

The experimental stimuli were 128 photographs that varied on dimensions of laterality (left or right), body part (hand or foot), and emotional valence (pain or no pain; 16 stimuli per category; Figure 1B) selected from the empathy-for-pain datasets used in Gu et al., (2010, 2013). These dimensions created eight categories: painful-left-hand, painful-left-foot, painful-right-hand, painful-right-foot, nonpainful-left-hand, nonpainful-left-foot, nonpainful-right-hand, and nonpainful-right-foot. Confirmative analyses showed that painful pictures received significantly higher pain ratings than nonpainful pictures [independent-sample *t* test: *t*_(126)_ = 48.99, *p* < 0.001, Cohen *d* = 8.66; pain: 3.41±0.35; no pain: 1.13±0.12]. The viewing angle for each stimulus was 5.97 × 5.32°.

**Figure1.**
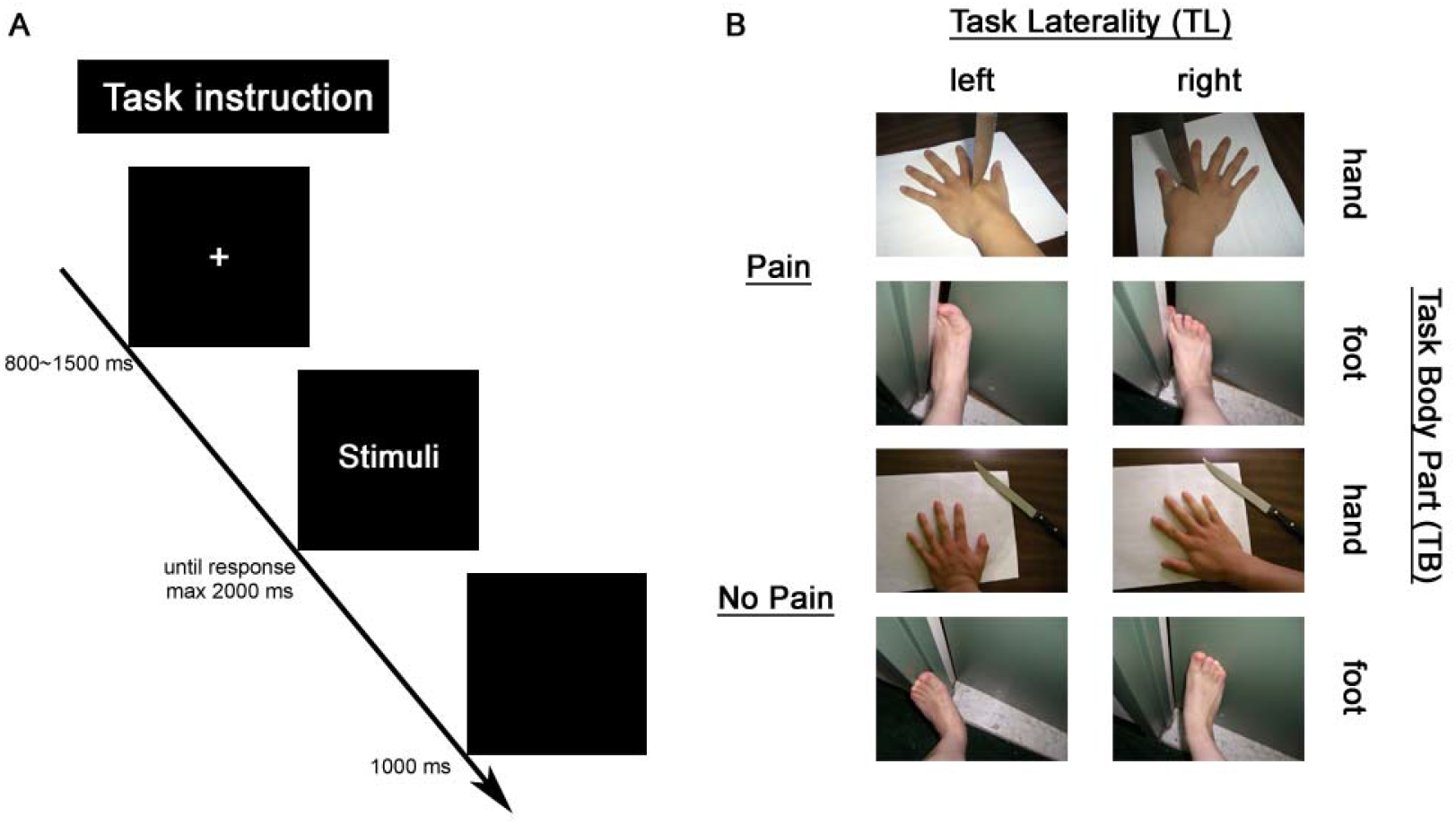
Experimental design. (A) Trial design of the Emotion-Cognition Integration (ECI) task. (B) examples of stimuli (Gu *et al.*, 2013).

**Figure2.**
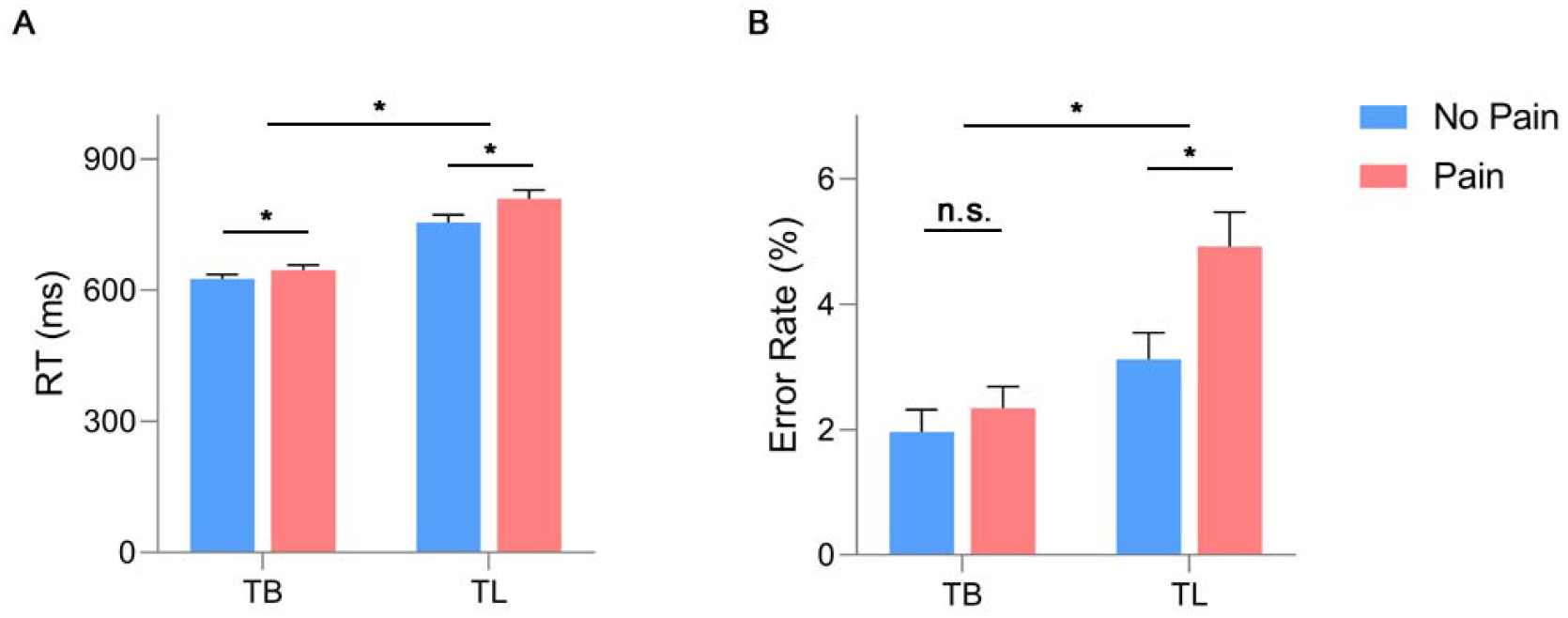
Behavioral results. (A) RT. (B) Error Rate. Abbreviations: RT, reaction time; TB, Task Body Part; TL, Task Laterality; * *p* < 0.05; n.s., *p* > 0.05.

**Figure3.**
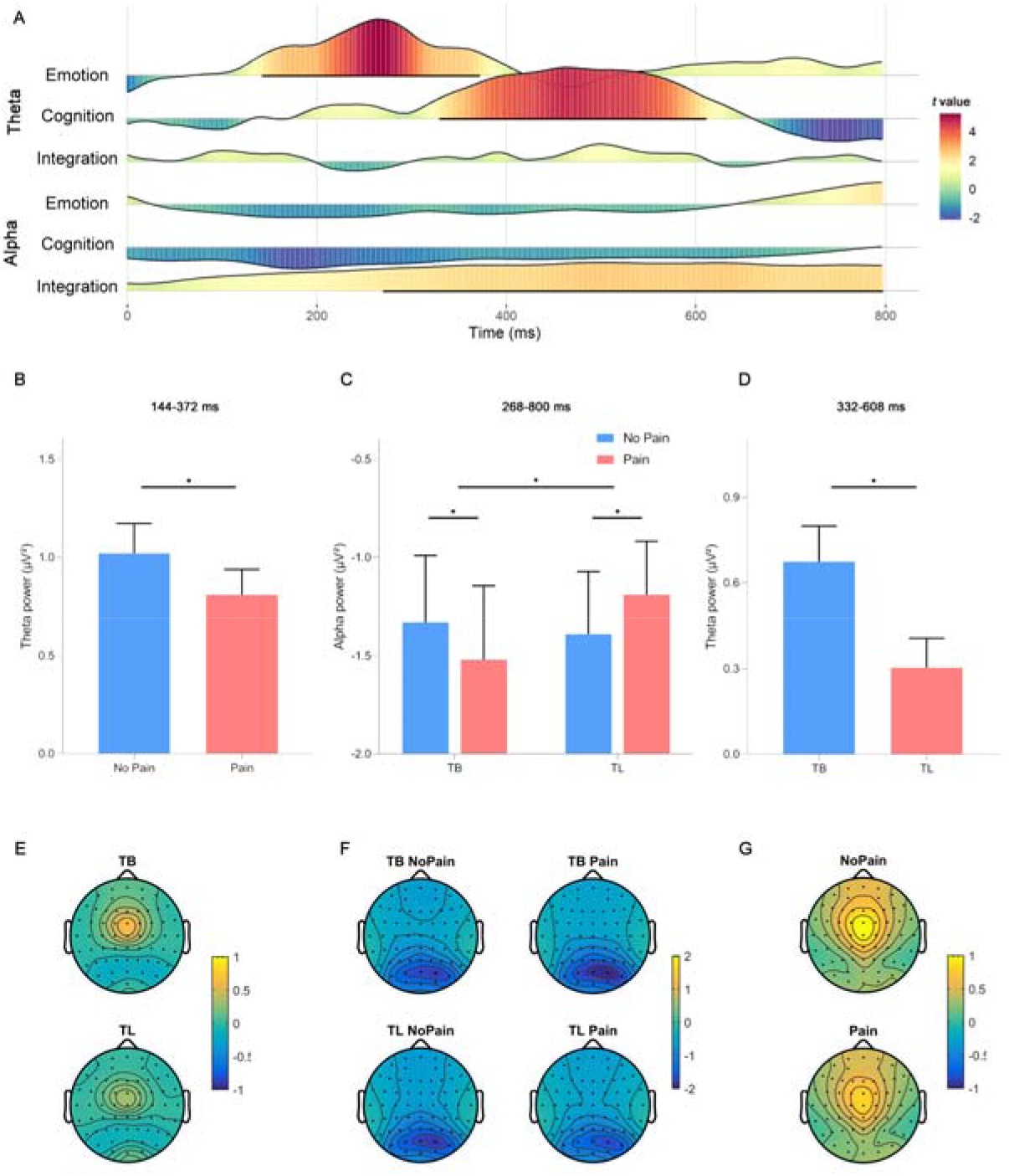
Oscillatory results. A) Time course of *t* values for each contrast at the onset of stimuli. Significant clusters were marked with the black solid line. B) The main effect of Emotion in the theta band from 144 to 372 ms; C) The interaction effect between Emotion and Cognition in the alpha band from 268 to 800 ms; D) The main effect of Cognition in the theta band from 332 to 608 ms. E) topographic maps for TB and TL conditions from 144 to 372 ms. F) topographic maps for each condition from 268 to 800 ms. G) topographic maps for Pain and NoPain conditions from 332 to 608 ms. Abbreviations: TB, Task Body Part; TL, Task Laterality; * *p* < 0.05.

### Task and Procedure

We used a modified ECI task (Figure 1A; Gu et al., 2013), which included two counterbalanced sessions: Task Laterality (TL) and Task Body Part (TB). The two tasks were identical except for the task instructions: In TL sessions, participants were asked to judge the laterality (left or right) of the presented body part. In TB sessions, participants were required to judge the body part (foot or hand) of the presented picture. As shown by Gu et al., (2013), TL was associated with higher cognitive demand than TB, inferred from significantly longer reaction times and higher error rates in TL than TB (mean reaction time (ms): TL = 1000.5, TB = 871; mean error rates (%): TL = 4.3, TB = 0.65). During each session, a total of 128 stimuli (64 for Pain and 64 for No Pain) were randomly presented. Each session consisted of two blocks, interleaved with a self-paced break. At the beginning of each trial, following a white fixation cross of a random duration (800~1500 ms), a picture was displayed in the center of the screen for 2000 ms maximally. Once they had identified the target (left or right in TL; hand or foot in TB) from the shown picture, participants pressed the button “j” with the left index finger or “k” with the right index finger. Specifically, TB required participants to press “j” if the photograph showed a hand and “k” if the photograph showed a foot. This response pattern was counterbalanced across participants. During TL, participants were instructed to press “j” if a left body part was shown and “k” if a right body part was shown. Finally, a blank screen lasted for 1000 ms. The experiment included four conditions with 256 trials (64 trials per condition). Participants completed several practice rounds until achieving at least 80% of accuracy before the formal experiment started. All experimental procedures were presented using E-prime 2.0 (Psychology Software Tools Inc. Pittsburgh, PA, USA).

### EEG Recording and Preprocessing

We recorded EEG data from a 64-electrode scalp cap according to the international 10-20 system (Brain Products, Munich, Germany), with the reference to the channel FCz. The electrooculogram (EOG; vertical) was recorded with electrodes placed below the right eye. Electrode impedances of EEG and EOG were maintained < 5 kΩ. All electrodes were amplified using a 0.01 online high-pass filter and continuously sampled at 1000 Hz per channel for offline analysis.

EEG data were preprocessed with EEGLAB 14.1.2b (Delorme and Makeig, 2004) in Matlab 2014b (MathWorks Inc). It comprised the following steps: 1) low-pass filtering of 30Hz; 2) resampling to 250Hz; 3) re-referencing offline to the average of the left and right mastoids (TP9 and TP10); 4) manually rejecting salient muscle sections and bad channels (if any); 5) Independent Component Analysis (ICA); 6) visually inspecting and rejecting artifact components (horizontal and vertical eye movements and muscle component); 7) interpolating bad channels (if any); 8) epoching from 1000 ms before to 1000 ms after stimuli onset; 9) baseline correction (−200 to 0 ms); 10) rejecting trials in which EEG voltages were out of range [−80, 80] μV.

### Behavioral statistics

We used SPSS 17.0 to perform statistical analyses, with the significance level at *p* = 0.05. Reaction times (RT) and Error Rates were subjected to a three-way ANOVA with Emotion (Pain/No Pain) and Cognition (TL/TB) as within-subject factors and with alexithymia group (HA/LA) as a between-subject factor.

### Oscillation statistics

Time-frequency distributions of each clean trial were computed by a short-time Fourier transform (STFT). With a hanning window of 250 ms and the method of detrend, we computed alpha and theta power for each point at the time domain (-1000 to 1000 ms; steps of 4 ms) and frequency domain (1 to 30 Hz; steps of 1Hz). We then performed baseline correction (−500 to −300 ms). Following the segmentation from 0 to 800 ms after stimuli onset, trials of time-frequency signals were included for statistical analyses only if the behavioral response was correct. Based on previous findings and our hypotheses, we focused on the frontal-central theta power (FCz; 4~7 Hz) and posterior alpha power (Pz; 8~12 Hz). Six contrasts were subjected to cluster-based permutation tests (paired *t* tests, two-tailed, 2000 times): 1) TB vs. TL in theta power; 2) No Pain vs. Pain in theta power; 3) (Pain – No Pain in TL) vs. (Pain – No Pain in TB) in theta power; 4) TB vs. TL in alpha power; 5) No Pain vs. Pain in alpha power; 6) (Pain – No Pain in TL) vs. (Pain – No Pain in TB) in alpha power. Bonferroni correction was used to control for multiple comparisons. Please note that the 3^rd^ and 6^th^ contrasts represented interaction effects between Emotion and Cognition. To check the direction for each effect, we then performed one-way repeated-measures ANOVA (between TL and TB or between Pain and No Pain) or two-way repeated-measures ANOVA with Cognition and Emotion as within-subject factors in the significant cluster (s). We did not take the group factor (HA/LA) into account here given that we did not find any group difference at the behavioral level (see Results section). Please note that a single peak with negatively skewed (skewness = −0.266 ± 0.306; kurtosis = −1.300 ± 0.604; Figure S1) was observed in the TAS-20 scores. Consequently, the data from the two groups (HA/LA) were combined for EEG statistical analyses.

## Results

### Behavioral results

For RT, the three-way ANOVA revealed a significant main effect of Emotion (*F*_(1,59)_ = 194.829, *p* < 0.001, η_p_^2^ = 0.768; Pain > No Pain; 95% CI: 31.980, 42. 684; Table 1) as well as a significant main effect of Cognition (*F*_(1,59)_ = 79.984, *p* < 0.001, η_p_^2^ = 0.575; TL > TB; 95% CI: 113.204, 178.461). We also observed a significant interaction effect between Emotion and Cognition (*F*_(1,59)_ = 45.939, *p* < 0.001, η_p_^2^ = 0.438). Simple effect analyses revealed longer RTs for Pain than No Pain in TL (*F*_(1,59)_ = 48.235, *p* < 0.001, η_p_^2^ = 0.450; 95% CI: 14.299, 25.873) relative to TB (*F*_(1,59)_ = 157.680, *p* < 0.001, η_p_^2^ = 0.728; 95% CI: 45.881, 63.275). No significant alexithymia-related effect was found (*ps* > 0.475). With regard to Error Rates, the three-way ANOVA showed significant main effects of Emotion and Cognition (For Emotion: *F*_(1,59)_ = 12.434, *p* = 0.001, η_p_^2^ = 0.174; Pain > No Pain; 95% CI: 0.005, 0.017; For Cognition: *F*_(1, 59)_ = 18.269, *p* < 0.001, η_p_^2^ = 0.236; TL > TB; 95% CI: 0.010, 0.027). Importantly, we found a significant interaction effect between Emotion and Cognition (*F*_(1, 59)_ = 6.530, *p* = 0.013, η_p_^2^ = 0.100). Based on simple effect analyses, Pain elicited higher Error Rates than No Pain in TL (*F*_(1, 59)_ = 16.274, *p* < 0.001, η_p_^2^ = 0.216; 95% CI: 0.009, 0.027), while no significant difference was found in TB (*F*_(1, 59)_ = 0.961, *p* = 0.331, η_p_^2^ = 0.016; 95% CI: −0.004, 0.012). No significant alexithymia-related effects were found (*ps* > 0.179).

**Table 1.** Behavioral and oscillatory responses in each experimental condition.

| **Emotion** | **Cognition** | **RT** | **Error Rate**<br>**(%)** | **Theta power**<br>**(144-372 ms)** | **Theta power**<br>**(332-608 ms)** | **Alpha power**<br>**(268-800 ms)** |
| --- | --- | --- | --- | --- | --- | --- |
| No Pain | Low | 626.02 (82.855) | 1.96 (2.814) | 1.06 (1.211) | 0.71 (1.076) | -1.33 (2.646) |
|  | High | 755.11 (138.72) | 3.12 (3.327) | 0.98 (1.231) | 0.30 (0.874) | -1.39 (2.476) |
| Pain | Low | 646.08 (92.06) | 2.34 (2.693) | 0.86 (1.235) | 0.64 (0.927) | -1.52 (2.927) |
|  | High | 809.80 (152.64) | 4.92 (4.308) | 0.76 (0.871) | 0.31 (0.773) | -1.19 (2.119) |

Abbreviations: Descriptive data were presented as mean (standard deviation). RT, reaction time.

### Oscillatory results

Among six contrasts, we found three significant clusters in theta and alpha bands following stimuli presentation: 1) a main effect of Emotion in theta band from 144 to 372 ms; 2) an interaction effect between Emotion and Cognition in alpha band from 268 to 800 ms; 3) a main effect of Cognition in theta band from 332-608 ms. Specifically, one-way ANOVAs showed that Pain evoked weaker theta power than No Pain from 144 to 372 ms (*F*_(1,60)_ = 17.630, *p* < 0.001, η_p_^2^ = 0.227; 95% CI: −0.313, −0.111) and TB evoked stronger theta power than TL (*F*_(1,60)_ = 18.718, *p* < 0.001, η_p_^2^ = 0.238; 95% CI: 0.199, 0.540). Particularly, a significant interaction between Emotion and Cognition from 268 to 800 ms (*F*_(1,60)_ = 6.036, *p* = 0.017, η_p_^2^ = 0.091) revealed increased alpha power in Pain relative to No Pain in TL (*F*_(1,60)_ = 5.692, *p* = 0.020, η_p_^2^ = 0.087; 95% CI: 0.032, 0.368) but decreased alpha power in Pain relative to No Pain in TB (*F*_(1,60)_ = 4.003, *p* = 0.050, η_p_^2^ = 0.063; 95% CI: 0.001, 0.382). Please note that the minimum number of trials in each condition was not less than 52.

## Discussion

Emotion-cognition interactions are of importance in determining human behavior.

Regarding their neural basis, electrophysiological mechanisms underlying their complex interrelationship (i.e., cooperation, competition or the combination) remain to be elucidated. Consistent with Gu et al., (2013) who examined the spatial organization of Emotion-Cognition Integration (ECI), our behavioral results showed increased RT and Error Rate in Pain relative to No Pain during TL as compared to TB, suggesting reciprocal competition and inhibition between emotion and cognition. Using high temporal resolution EEG recording, we found that 1) emotion processing occurred first in the theta band from 144 to 372 ms; 2) ECI took place from 268 ms and lasted to 800 ms in the alpha band; 3) the theta band also reflected cognitive processing, which occurred from 332 to 608 ms. These findings shed light on the temporal organization of brain oscillations underlying ECI processes, with emotion processing occurring first and cognition processing occurring later in the theta band. ECI took place in the alpha band.

Theta, rather than alpha, power reflected emotion and cognition processing at different stages, while alpha, but not theta, power was responsible for integrating computation between emotion and cognition, indicative of a functional dissociation between the theta and alpha oscillations in ECI processes. Theta oscillations are involved in fundamental functions across species (Knyazev, 2007). In addition to emotion processing and cognitive encoding (Knyazev, 2007; Kamarajan *et al.*, 2008; Knyazev *et al.*, 2009; Cavanagh and Frank, 2014), theta power has been linked to cognitive regulation of negative emotion (Ertl *et al.*, 2013). However, inconsistent with our prediction, we did not find any significant integrative cluster in the theta band, which may suggest that ECI is associated with different neural mechanisms than cognitive regulation of negative emotion. Notably, the prefrontal cortex and the amygdala are at the core of emotion regulation (Kohn *et al.*, 2014; Xu *et al.*, 2020; Wang, Goerlich, *et al.*, 2021), while the anterior insula may be the core brain area underlying ECI (Gu *et al.*, 2013; Luo *et al.*, 2014). Future studies would benefit from investigating differences in alpha and theta oscillations between emotion regulation and ECI. Although many studies have linked increased theta power to higher cognitive demand (Nigbur *et al.*, 2011; Cavanagh and Frank, 2014), some studies have reported the reverse pattern (Puma *et al.*, 2018; Kramer, 2020). It has been proposed that task difficulties moderate the relationship between theta power and cognitive demand in an inverted U-shape pattern, with theta oscillations decreasing for simple tasks, reaching highest levels for medium task difficulty and decreasing again with high task demand (Puma et al., 2018). We found decreased theta power in tasks with low vs. high cognitive demand. One explanation might be that the current task was relatively easy for the participants, as suggested by the high accuracy in the current study. Among ECI processes, emotion processing took place first, from 144 to 372 ms. This result supports the dual-system theory distinguishing between a fast affective system and a slow cognitive system in the human brain (Kahneman, 2003; Kuo *et al.*, 2009).

Importantly, ECI was subserved by activity in the alpha band from 268 to 800 ms. Alpha oscillations have long been considered dominant in the human brain (Başar, 2012), with mounting evidence supporting an inhibitory function of alpha oscillations in information processing (Klimesch *et al.*, 2007; Jensen and Mazaheri, 2010). Increased alpha power may reflect inhibition, while decreased alpha power may denote a release from inhibition (or facilitation; Klimesch, 2012). These roles of alpha activity have been considered to be part and parcel of integrative processes underlying several functions, including attention, working memory, and emotion (Başar, 2012; Klimesch, 2012; Van Diepen *et al.*, 2019).

For example, alpha power has been shown to suppress cognitive control of emotional distractions (Murphy *et al.*, 2020). The current results of increased alpha power in pain (vs. neutral) processing during tasks with high cognitive demand, in line with our behavioral findings, may thus be indicative of mutual inhibition between emotion and cognition to compete for limited mental resources.

With regard to the time scale, integrative operation in the alpha band occurred between 268 and 800 ms (note the segmentation of each trial from 0 to 800 ms post-stimulus onset and ECI most likely continuing after 800 ms). The relatively early and lasting integrative processing between emotion and cognition suggests an indispensable role of integrative computation, challenging the traditional view of emotion and cognition being independent systems (Goldman-Rakic, 1996; LeDoux, 2000; Ochsner and Gross, 2005; Pessoa, 2008; Liu *et al.*, 2009). In sum, our findings contribute to developing a temporal architecture underlying ECI during an empathy-for-pain paradigm. Future studies would benefit from obtaining a full picture of ECI processes by simultaneous EEG-fMRI or Magnetoencephalography recordings or intracranial EEG recording. An advantage for intracranial EEG recording is that the high-gamma band can be reliably accessed given its crucial role in information integration (Zhang and Yartsev, 2019; Wang *et al.*, 2020).

We also found decreased alpha power in pain (vs. neutral) processing during low cognitive demand (body part judgement) following the interaction between emotion and cognition in the alpha band. Based on the facilitation-inhibition hypothesis regarding alpha oscillations (Klimesch, 2012; Van Diepen *et al.*, 2019), this result suggests mutual facilitation between emotion and cognition. Although such a facilitating effect was indeed observed in RT in the previous study (Gu *et al.*, 2013), it was not observed in the current behavioral results. Therefore, interpretations regarding this result should be made with caution. Please note that we did not find any significant interaction effect between emotion and cognition in pain for empathy-related and cognition-related event-related potentials (ERPs; see Figure S2; Nieuwenhuis et al., 2011; Coll, 2018). Compared to our oscillation results, the ERP results may suggest that phase-locked analysis is not sensitive to ECI signals.

No significant alexithymia-related effect was observed in the current study, despite its association with impairments in emotional processing (van der Velde *et al.*, 2013; Wang, Chen, *et al.*, 2021). One possible explanation may be that the stimuli used here merely displayed isolated parts of the human body and thus lacked social-emotional features (Britton et al., 2006; Powers et al., 2013). In a previous study, we reported social-specific impairment for negative emotion processing in alexithymia (Wang, S Goerlich, *et al.*, 2021). Future studies could examine the integrative processing between emotion and cognition in relation to alexithymia with socially relevant materials.

Several limitations of the present study should be mentioned. First, we examined the temporal organization of ECI only from electrophysiological activity (e.g., oscillation) profiles. It has been proposed that complex cognitive-emotional behaviors are organized in dynamic brain networks (Pessoa, 2008). Future studies should aim to explore neural mechanisms of ECI using functional connectivity. Second, we used photographs that varied only between painful and nonpainful levels in the current study, because empathic pain has been considered an excellent candidate to probe ECI (Gu *et al.*, 2013). It is thus not clear whether the current conclusions can be extrapolated to other emotions. Future studies would thus benefit from the examination of ECI processing in a variety of emotional categories.

To conclude, this study shows characteristics of oscillatory dynamics underlying ECI, with emotion processing occurring first and cognition processing occurring later in the theta band and ECI taking place in the alpha band. In support of the dual-system theory of decision making (Kahneman, 2003; Kuo *et al.*, 2009), our results are indicative of fast processing of emotional information, slower cognitive processing, and mutual inhibition between emotion and cognition to compete for limited mental resources. These findings shed light on the role of neural oscillations on the interplay between emotion and cognition in the human brain.

## Data available statement

The data are available from the corresponding author P. Xu, upon reasonable request.

