## Supplement for "Oscillatory dynamics underlying emotion-cognition integration: differential role of theta and alpha oscillations"

**ERP results**

In addition to oscillatory results, we also analyzed empathic pain-related and cognitive-related ERPs. For N1 (Fz, FCz, and Cz; 110~150 ms), N2 (Fz, FCz, and Cz; 220~280 ms), P3 (POz, PO3, and PO4; 330~420 ms), and LPP (Pz, P1, and P2; 500~800 ms), two-way repeated measures ANOVAs were performed with Emotion (Pain/No Pain) and Cognition (TB/TL) as within-subject factors. For N1, we observed a marginally significant main effect of Emotion [$F_{(1,60)} = 3.881$, $p = 0.053$, $\eta_p^2 = 0.061$; Pain > No Pain; 95% CI: -0.004, 0.548]. No other significant effect was found ( both $p > 0.438$). Regarding N2, we observed a significant main effect of Emotion [$F_{(1,60)} = 85.683$, $p < 0.001$, $\eta_p^2 = 0.588$; Pain > No Pain; 95% CI: 1.421, 2.204]. No other significant effect was found ( both $p > 0.506$). With regard to P3, we found significant main effects of both Emotion [$F_{(1,60)} = 4.162$, $p = 0.046$, $\eta_p^2 = 0.065$; Pain < No Pain; 95% CI: -0.573, -0.006] and Cognition [$F_{(1,60)} = 5.443$, $p = 0.023$, $\eta_p^2 = 0.083$; TL < TB; 95% CI: -1.049, -0.080]. No significant interaction effect between Emotion and Cognition was found [$F_{(1,60)} = 1.470$, $p = 0.230$, $\eta_p^2 = 0.024$] . For LPP, we observed a significant main effect of Emotion [$F_{(1,60)} = 66.676$, $p < 0.001$, $\eta_p^2 = 0.526$; Pain > No Pain; 95% CI: 1.159, 1.911]. No other significant effect was found ( both $p > 0.147$). In sum, we did not find any interaction effect between Emotion and Cognition in N1, N2, P3, and LPP except main effects of Emotion and Cognition (Figure S2).

**Group analysis on the alpha-band cluster**

To check whether there was a group difference in integrative oscillations (alpha power), we performed a three-way repeated-measures ANOVA with Cognition and Emotion as within-subject factors and with Group as a between-subject factor in the significant alpha cluster. No significant group-related effect was found (*ps* > 0.130).

**Exploratory correlational analysis of ECI effects in reaction times, error rates, and alpha oscillations with facets of alexithymia**

To check whether there were associations between ECI effects and facets of alexithymia, we performed Spearman correlations. Here ECI effects were defined as [(Pain – No Pain) in TL – (Pain – No Pain) in TB] in reaction times, error rates, and alpha oscillations, respectively. Alexithymia facets included difficulty identifying feelings, difficulty describing feelings, externally oriented thinking of the TAS-20, verbalizing, fantasizing, successful identifying, unsuccessful identifying, emotionalizing, and analyzing subscale of BVAQ, as well as second-order cognitive and affective factors of BVAQ. Results showed a significant positive correlation between difficulty describing feelings of the TAS-20 and alpha oscillations (rho = 0.288, *p* = 0.025) and a significant negative correlation between the analyzing subscale of BVAQ and error rates (rho = -0.257, *p* = 0.046). Please note that higher scores in the TAS-20 represent higher levels of alexithymia, while lower scores in the BVAQ represent higher levels of alexithymia. These correlational results may suggest that individuals with difficulty in describing and analyzing feelings have a stronger ability of ECI. However, interpretations should be cautious given the nature of exploratory analyzes without multiple comparison corrections.

**Figure S1. Distribution of TAS-20 scores.**

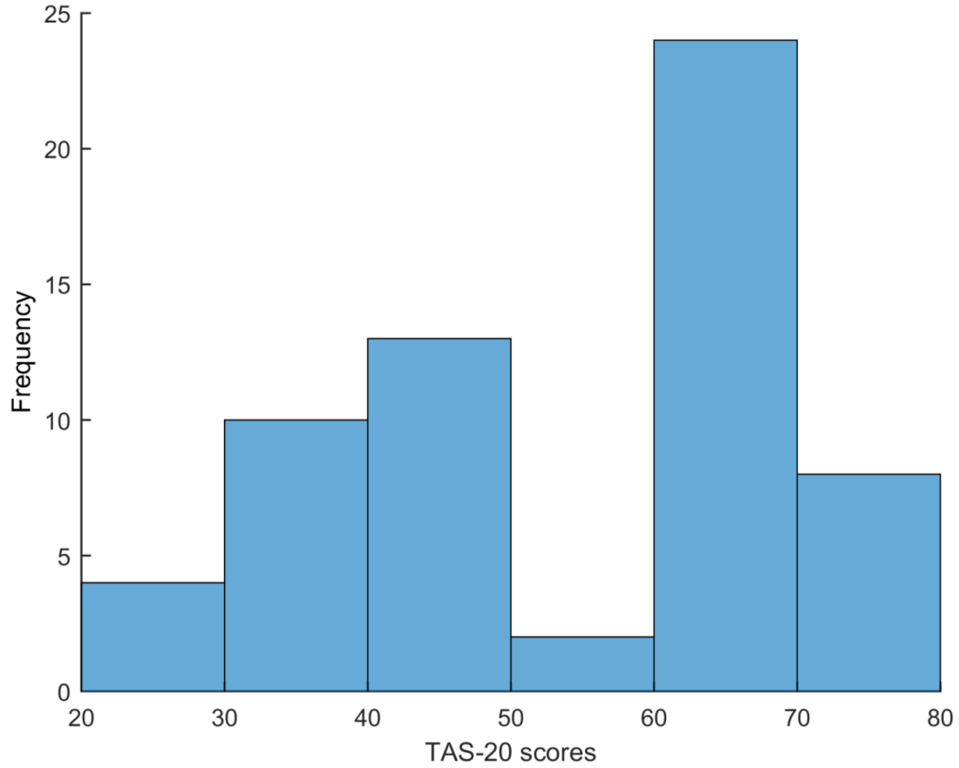

**Figure S2. ERP results.** Time course in FCz (A), POz (B), and Pz (C) electrodes. (D) topographic map within N1 window for NoPain and Pain conditions. (E) topographic map within N2 window for NoPain and Pain conditions. (F) topographic map within P3 window for NoPain and Pain conditions. (G) topographic map within P3 window for NoPain and Low and High cognitive demand conditions. (H) topographic map within LPP window for NoPain and Pain conditions. Statistical results of main effects of Emotion for N1 (I), N2 (J), P3 (K), LPP (M). (L) Statistical results of main effects of Cognition in P3.

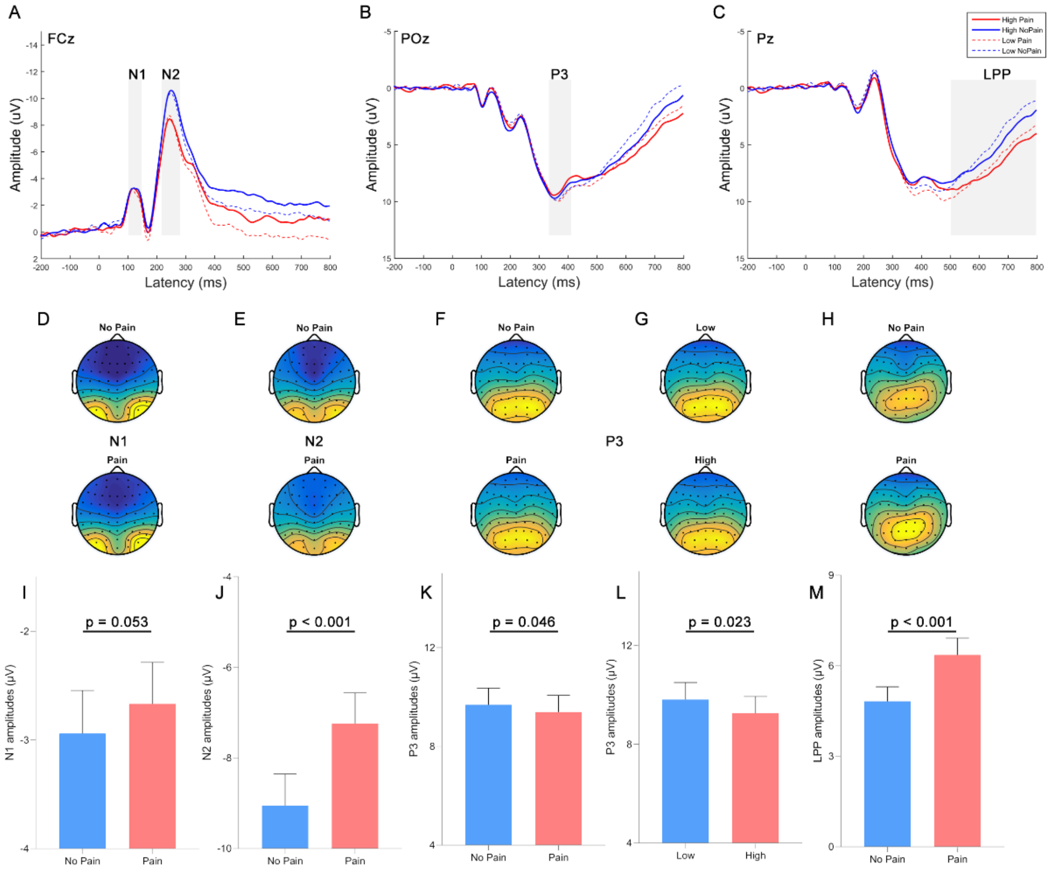

**Table S1.** Demographics and questionnaire scores of participants.

| | **HA (32; 16 females)** | | **LA (29; 16 females)** | | **t** | **p** |
|---|---|---|---|---|---|---|
| | Mean (SD) | [min, max] | Mean (SD) | [min, max] | | |
| Age | 20.06 (1.78) | [17, 24] | 20.97 (2.60) | [18, 28] | -1.57 | 0.123 |
| TAS-20 | 66.56 (4.74) | [61, 77] | 39.07 (7.32) | [25, 51] | 17.22 | <0.001 |
| DIF | 24.34 (3.31) | [17, 31] | 11.66 (4.11) | [7, 21] | 13.34 | <0.001 |
| DDF | 18.25 (2.27) | [14, 22] | 9.41 (1.99) | [6, 14] | 16.08 | <0.001 |
| EOT | 23.97 (2.92) | [18, 31] | 18.00 (4.16) | [10, 23] | 6.42 | <0.001 |
| BAI | 32.88 (7.31) | [22, 45] | 25.72 (4.70) | [21, 37] | 4.59 | <0.001 |
| BDI | 14.44 (8.93) | [1, 38] | 4.38 (5.06) | [0, 19] | 5.48 | <0.001 |
| ASQ | 122.72 (8.94) | [102, 139] | 111.76 (10.35) | [87, 128] | 4.44 | <0.001 |
| BVAQ | 110.81 (13.29) | [64, 130] | 134.55 (14.52) | [110, 169] | -6.67 | <0.001 |
| BVAQ_V | 17.41 (3.91) | [10, 26] | 27.07 (4.34) | [19, 34] | -9.15 | <0.001 |
| BVAQ_F | 29.97 (6.06) | [12, 40] | 29.55 (5.95) | [15, 40] | 0.27 | 0.787 |
| BVAQ_SI | 9.28 (2.28) | [4, 13] | 12.38 (2.09) | [8, 15] | -5.51 | <0.001 |
| BVAQ_UI | 13.44 (2.61) | [8, 19] | 20.86 (3.06) | [15, 25] | -10.22 | <0.001 |
| BVAQ_E | 20.50 (3.65) | [8, 26] | 20.59 (3.34) | [15, 27] | -0.10 | 0.924 |
| BVAQ_A | 20.22 (4.15) | [10, 29] | 24.10 (3.32) | [17, 29] | -4.01 | <0.001 |
| Affective | 50.47 (7.69) | [29, 64] | 50.13 (8.37) | [31, 67] | 0.16 | 0.873 |
| Cognitive | 60.34 (8.50) | [35, 75] | 84.41 (10.22) | [66, 102] | -10.04 | <0.001 |

Note: HA, individuals with high alexithymia; NonALEX, individuals with low alexithymia; TAS-20, 20-item Toronto Alexithymia Scale; DIF, difficulty identifying feelings; DDF, difficulty describing feelings; EOT, externally oriented thinking; BAI, Beck Anxiety Inventory; BDI, Beck Depression Inventory. ASQ, Autism Spectrum Quotient; BVAQ, Bermond-Vorst Alexithymia Questionnaire; BVAQ_V, verbalizing subscale of BVAQ; BVAQ_F, fantasizing subscale of BVAQ; BVAQ_SI, successful identifying subscale of BVAQ; BVAQ_UI, unsuccessful identifying subscale of BVAQ; BVAQ_E, emotionalizing subscale of BVAQ; BVAQ_A, analyzing subscale of BVAQ; Affective, affective component of BVAQ; Cognitive, cognitive component of BVAQ.

**Table S2.** ERP amplitudes in each experimental condition.

| Emotion | Cognition | N1 | N2 | P3 | LPP |
|---|---|---|---|---|---|
| No Pain | Low | -2.95 (3.417) | -8.96 (5.586) | 9.89 (5.310) | 4.51 (3.783) |
| | High | -2.92 (2.940) | -9.14 (5.577) | 9.48 (5.399) | 5.13 (4.086) |
| Pain | Low | -2.58 (2.992) | -7.25 (5.628) | 9.76 (5.391) | 6.23 (4.501) |
| | High | -2.76 (3.214) | -7.23 (5.171) | 9.03 (5.281) | 6.48 (4.713) |

Abbreviations: Descriptive data are presented as mean (standard deviation).
